# Macrophages promote collagen deposition through circadian regulation of fibroblasts

**DOI:** 10.1101/2025.02.25.640115

**Authors:** Katherine Lowles, Marie F.A. Cutiongco, Joshua J. Hughes, Shiyang Li, John Knox, Madeleine Coy, Wei-Hsiang Lin, Andrew S. MacDonald, Karl E. Kadler, Qing Jun Meng, Tracy Hussell, Joan Chang

## Abstract

Collagen deposition in fibroblasts, the primary collagen-producing cells, is regulated by both macrophages and the circadian rhythm, although how these regulatory processes interact with one another is unknown. Here, we reveal that macrophage-fibroblast interaction enhances collagen deposition which requires a functional circadian rhythm. This response is also dependent on macrophage polarisation status, where M0 (naïve) and M1 (pro-inflammatory) macrophages require direct cell-cell contact; whereas M2 (anti-inflammatory) macrophages utilise an additional mechanism through secreted soluble factors that strengthen circadian rhythms and significantly increase collagen deposition in fibroblasts, independent of cell-cell contact. Using mass spectrometry proteomics analysis, we identified PDGFA as a key factor in induction of circadian amplitude in fibroblasts. Crucially, macrophages lacking PDGFA showed diminished ability to modulate fibroblast circadian rhythms and collagen production. Further, in fibroblasts with impaired circadian rhythms, neither macrophages nor macrophage-secreted factors are capable of inducing collagen deposition. These results confirmed that the collagen secretory pathway is also under circadian clock control in lung fibroblasts, and demonstrated that macrophages influence collagen deposition via circadian clock mediated mechanisms. In conclusion, our findings highlight M2-derived PDGFA as a key macrophage-derived factor in extracellular matrix remodelling.

## Introduction

Type-I collagen (collagen) is one of the most abundant extracellular matrix (ECM) proteins, and its homeostatic control is crucial for tissue health. Dysregulation of collagen leads to conditions such as fibrosis and chronic wounds^1^. The key cells involved in collagen regulation are fibroblasts and macrophages. Fibroblasts are the predominant collagen producing cells, while macrophages play a regulatory role by secreting cytokines and growth factors that influence collagen synthesis in fibroblasts^2,3^. Intriguingly, macrophages have also been reported to directly deposit collagen at wound sites, suggesting additional regulatory roles in the ECM. However, it is unclear if this deposition is also observed under homeostatic conditions^4^.

Recent studies have demonstrated that in collagen-rich tissues, such as tendons, there is a daily surge of collagen production. Certain genes involved in collagen biosynthesis - particularly those that regulate trafficking through the secretory pathway - are controlled by the circadian clock^5^. Around 50% of mammalian genes are rhythmically expressed with any 24-hour cycle, thus maintaining appropriate rhythms in these clock-controlled genes (CCGs) is critical for homeostasis and health^6,7^. Uncoupling CCGs from the circadian clock results in impaired wound healing and contributes to a host of ECM-related diseases, including lung fibrosis^8,9^. It is crucial to understand the mechanisms that regulate rhythmic collagen secretion, in order to determine how these processes are disrupted under non-homeostatic conditions such as during injury responses, thus guiding the development of therapeutic strategies to restore a healthy, balanced state.

Macrophages play a pivotal role in tissue remodelling and repair by influencing fibroblast activity and collagen deposition^10^. Through direct cell-cell interactions and the secretion of soluble factors, macrophages modulate fibroblast behaviour, including their capacity to produce and organise collagen within the ECM^11^. This crosstalk is essential for maintaining tissue homeostasis and responding to injury, and disruption may drive fibrotic responses. Importantly, macrophage polarisation states—such as the pro-inflammatory M1 and pro-reparative M2 phenotypes— differentially regulate fibroblast activity, suggesting that macrophages fine-tune collagen dynamics depending on the physiological or pathological context^12,13^. As macrophages are known to localise to sites of high ECM/collagen deposition and remodelling together with fibroblasts, it is likely that macrophages may influence the fibroblast circadian clock, to synchronise and enhance rhythmic collagen production.

We hypothesised that macrophages influence collagen deposition in fibroblasts by modulating the fibroblast circadian clock. To test this, we assessed the impact of macrophages on collagen deposition using immunofluorescence (IF) imaging and investigated circadian rhythms in fibroblasts using the *Per2*-Luc circadian reporter (where luciferase is used as a readout for the circadian core clock protein Per2). Here, we showed for the first time that macrophages, regardless of polarisation status (M0 – naïve; M1 – LPS and IFN-γ stimulated; M2 – IL-4 and IL-13 stimulated), enhanced circadian rhythms of lung fibroblasts. Furthermore, we observed an increased collagen deposition in co-cultures. Intriguingly, M0 and M1 macrophages require cell-cell contact with fibroblasts to exert their influence, whereas secreted factors from M2 macrophages alone are sufficient to drive collagen fibrillogenesis and circadian rhythm induction. Using mass spectrometry (MS) proteomics analysis, we identified growth-factor PDGFA as a key M2 macrophage-secreted factor influencing fibroblast function. Notably, PDGFA alone can induce a circadian rhythm but is insufficient to promote collagen fibrillogenesis. Importantly, knocking out PDGFA in macrophages abrogated their ability to promote collagen deposition and enhance circadian rhythms in fibroblasts. Additionally, the macrophage-induced effects were dependent on a functional circadian rhythm in fibroblasts. Taken together, these results demonstrate that macrophages promote collagen deposition in fibroblasts by inducing circadian rhythms, with M2-derived PDGFA playing a key role in this process.

## Results

### Macrophages induce robust circadian rhythms in fibroblasts via myeloid-specific mechanisms

The interaction between macrophages and the circadian clock in fibroblasts was explored using fibroblasts isolated from the *Per2*-Luc (clock protein Period2 fused with luciferase protein) mice, where luminescence is released in the presence of luciferin and represents a direct readout of the abundance of Per2 protein; this provides real-time tracking of the circadian rhythm in fibroblasts^23^. Human cell-line THP-1 monocytes and macrophages, as well as primary AMs were added to *Per2*-Luc fibroblasts approximately 24 hours after bioluminescence recording began.

Untreated *Per2*-Luc lung fibroblasts (black trace, Figure 1A) showed an initial small peak that represented the overall sum of endogenous circadian rhythms of lung fibroblasts in culture, which are synchronised at low levels, likely through mechanical force exerted during media change and placement of culture dishes in the LumiCycle apparatus. This synchronisation rapidly diminished, reflecting desynchronisation caused by weak intercellular clock-clock coupling and cell division^24^.

**Figure 1.**
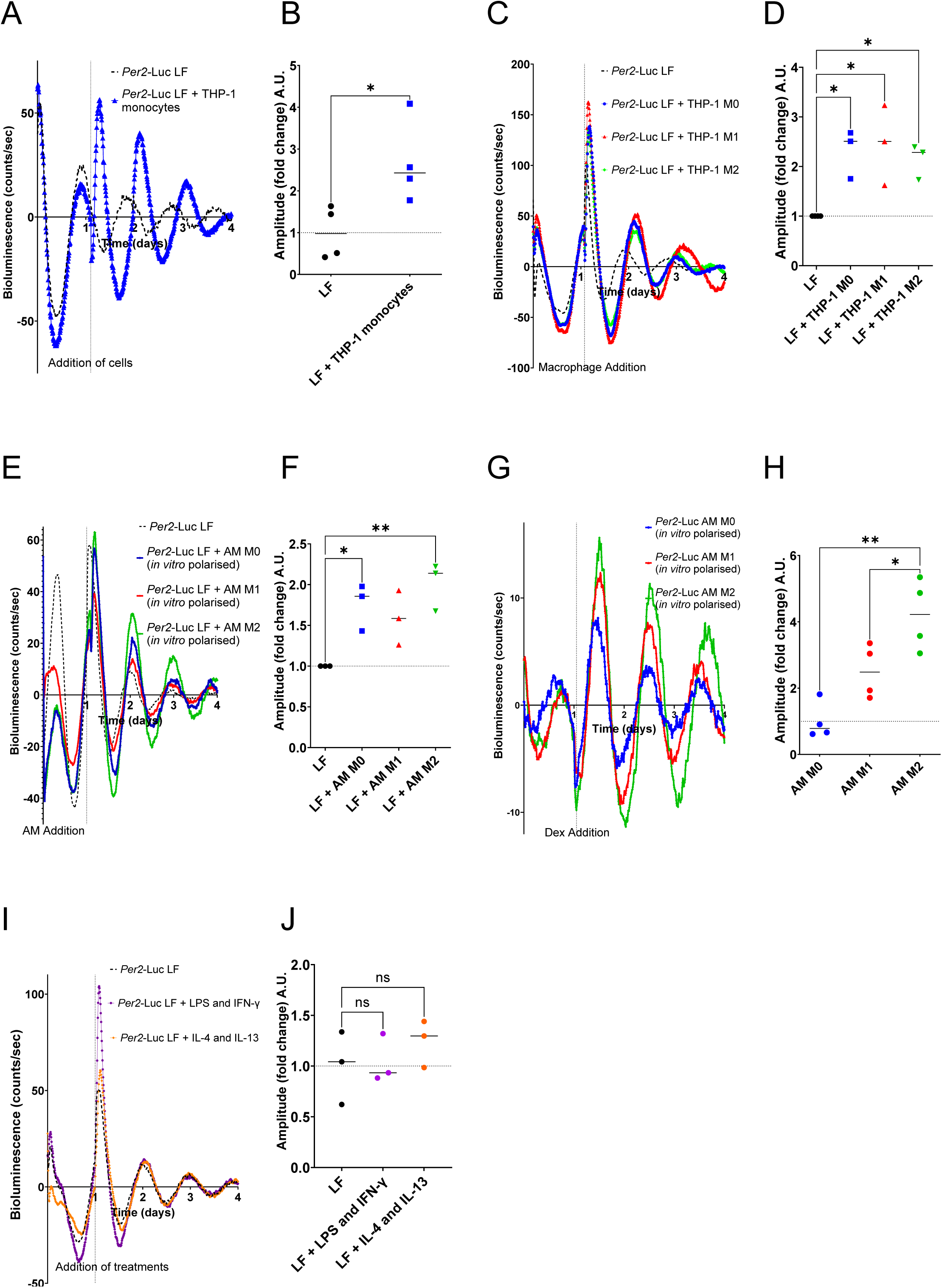
Bioluminescence Readings of *Per2*-Luc Lung Fibroblasts with Polarised THP-1 Macrophages. (A) Addition of THP-1 monocytes induced a robust circadian rhythm in fibroblasts. Representative of N= 4. (B) Amplitude fold change was shown to be significant following monocyte addition. N = 4 biological repeats, *p<0.05, one-way ANOVA. (C) Addition of M0, M1, and M2 polarised THP-1 macrophages induced robust rhythms in *Per2*-Luc fibroblasts, significantly increasing bioluminescence amplitude. Representative of N= 3. (D) Amplitude fold change showed significant increases with M0, M1 and M2, compared to LF alone. N = 3 biological repeats, *p<0.05, one-way ANOVA. (E) Addition of *in vitro* polarised primary murine AMs (M0 and M2) also induced robust rhythms in fibroblasts, with M2 having the greatest effect. Representative of N= 3. (F) Amplitude fold change showed significant increases with M0 and M2. N = 3 biological repeats, **p<0.01, one-way ANOVA. (G) M2-polarised *Per2*-Luc macrophages sustained the circadian rhythm with the highest amplitude following dexamethasone addition. Representative of N= 3. (H) Amplitude fold change analysis revealed that M2 macrophages significantly enhanced circadian amplitude compared to M0 and M1 macrophages. N = 4 biological repeats, *p<0.05, **p<0.05, one-way ANOVA. (I) M1- and M2-polarising factors were added to lung. Neither LPS and IFN-□, nor IL-4 and IL-13 (M1 and M2 polarising factors, respectively) increased circadian amplitude. Representative of N=3. (J) Quantification of amplitude fold change showed there was no significant increase in fibroblast circadian amplitude following addition of M1- and M2-polarising factors. N=3 biological repeats, one-way ANOVA.

Addition of THP-1 monocytes (blue trace, Figure 1A) after ∼24 hours resulted in a sustained and robust circadian rhythm in fibroblasts for over 2 days. Quantification of amplitude changes (Figure 1B) demonstrated that macrophages significantly enhanced circadian rhythm amplitude. In contrast, the addition of the epithelial Eph4 cells did not significantly increase amplitude (Supplementary Figure 1A, B), suggesting a macrophage-specific effect.

To confirm if THP-1 macrophages can also induce a rhythm in fibroblasts, we differentiated them first to M0 using PMA, followed by polarisation to M1 (pro-inflammatory, defined as 48-hour treatment with LPS and IFN-γ in this study), or M2 (anti-inflammatory, defined as 48-hour treatment with IL-4 and IL-13 in this study) states. This allowed us to determine whether effects on fibroblast circadian rhythms are dependent on macrophage polarisation status. Polarisation was confirmed through qPCR analysis of classical M1 (CD64) and M2 (TGF-β) gene markers (Supplementary Figure 1C, D); here, we found that M0, M1, and M2 macrophages all significantly increased circadian rhythm amplitude in fibroblasts (Figure 1C, D), with no significant differences between polarisation states.

We then asked if the circadian status of the macrophages influence their synchronisation ability. Macrophages were synchronised with dexamethasone, a glucocorticoid known to synchronise circadian rhythm, for 1 hour or 24 hours prior to addition to *Per2*-Luc fibroblasts^25^. The synchronisation status of macrophages did not alter their rhythm-enhancing effects on fibroblasts; unsynchronised, 1-hour synchronised, and 24-hour synchronised macrophages all induced significant amplitude increases (Supplementary Figure 1E, F).

To confirm that induction of circadian rhythm in lung fibroblasts was not due to cross-species reactivity, we used primary murine alveolar macrophages AMs isolated from lungs of wild-type (WT) mice (i.e. no luciferase activity). Macrophages were then polarised *in vitro,* as confirmed by qPCR analyses (Supplementary Figure 1G, H), and then added to *Per2*-Luc lung fibroblasts. Here, M2-polarised AMs induced the most robust rhythm in fibroblasts, followed by M0, while M1 AMs had minimal effect (Figure 1E, F). Similarly, *in vivo* polarised primary murine AMs were also assessed for their circadian-inducing effects. While M0, M1 and M2 macrophages all increased circadian amplitude in fibroblasts, only M2 *in vivo* polarised macrophages had a significant effect (Supplementary Figure 1I, J).

To evaluate macrophage-autonomous rhythms, AMs from *Per2*-Luc transgenic mice were polarised and synchronised with dexamethasone. All AMs showed a synchronisation response in their circadian rhythm; although amplitude measurements showed that, compared to M0 and M1, the amplitude increase in M2 AMs was significant. This suggests that M2 AMs inherently have the highest circadian amplitude and most sustained rhythm amongst the polarisation statuses (Figure 1G, H). Importantly, addition of M1-polarising factors (IFN-γ, LPS) or M2-polarising factors (IL-4, IL-13) to fibroblasts failed to enhance circadian amplitude, indicating it is a myeloid cell effect, and not a cytokine effect (Figure 1I, J).

Furthermore, THP-1 cells were transduced with a PER2-Luc2 plasmid in order to measure the circadian rhythms of M0 macrophages following the addition of WT fibroblasts (Supplementary Figure 1K). Fibroblasts did not increase the circadian amplitude in M0 macrophages (Supplementary Figure 1L). Taken together, these results demonstrated that macrophages can induce a circadian rhythm in fibroblasts, and that this circadian induction effect is not reciprocal.

### Macrophage-secreted factors are responsible for the circadian- and fibril-induction effects on fibroblasts□

To investigate if macrophages influence circadian amplitude through the secretion of soluble factors, serum-free CM from polarised macrophages was collected and added to *Per2*-Luc fibroblasts. Both THP-1 M0 and M2 CM increased circadian amplitude in the fibroblasts (Figure 2A). However, only CM from THP-1 M2 macrophages resulted in a statistically significant amplitude enhancement (Figure 2B).

**Figure 2.**
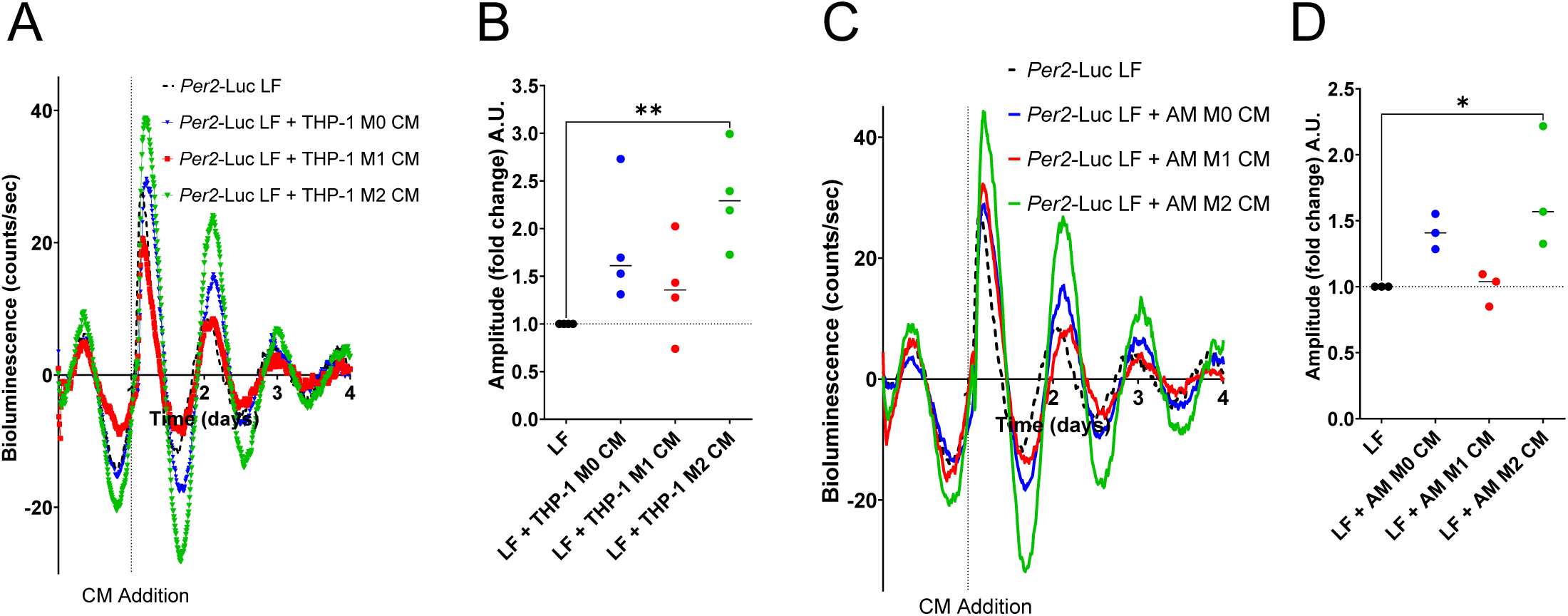
Bioluminescence Readings of *Per2*-Luc Lung Fibroblasts with Polarised THP-1 Macrophage Conditioned Media. (A) CM from polarised THP-1 macrophages was added to *Per2*-Luc lung fibroblasts. M2 macrophage CM induced a robust circadian rhythm in fibroblasts. Representative of N= 4. (B) Amplitude fold change analysis showed that while M0 CM and M2 CM increased circadian amplitude, only the increase with M2 CM was statistically significant compared to fibroblasts without CM. N = 4 biological repeats, **p<0.01, one-way ANOVA. (C) CM from M2-polarised AMs, and to a lesser extent M0 AMs, enhanced circadian amplitude in *Per2*-Luc lung fibroblasts. Representative of N= 3. (D) M2 CM significantly increased circadian amplitude compared to untreated *Per2*-Luc fibroblasts. N = 3 biological repeats, *p<0.05, one-way ANOVA.

Similarly, CM from *in vitro* polarised primary AMs was also added to *Per2*-Luc fibroblasts. CM from AM M0, M1, and M2 macrophages all promoted an increase in circadian amplitude (Figure 2C). A significant amplitude increase was observed exclusively with AM M2 macrophage CM (Figure 2D). These findings suggest that macrophage-derived soluble factors, particularly those released by M2-polarised macrophages, play a key role in enhancing circadian amplitude in fibroblasts.0

### Macrophages increase collagen deposition when co-cultured with fibroblasts

To investigate the impact of macrophages on collagen deposition in fibroblasts, *in vitro* co-culture experiments were performed, followed by immunofluorescence (IF) analysis to visualise and quantify changes in fibroblast-derived collagen. Co-cultures of polarised THP-1 macrophages and fibroblasts were seeded at a 1:5 ratio, as this is similar to the ratio between macrophages and fibroblasts in human lungs^26–28^. Collagen deposition was visualised using confocal microscopy (Figure 3A), and collagen area was quantified in these cultures with results normalised to nuclei numbers (Figure 3B). Lung fibroblasts co-cultured with M0-, M1-, or M2-polarised THP-1 macrophages all deposited significantly more collagen fibrils compared to fibroblasts cultured alone. Fibronectin staining in fibroblast-macrophage co-cultures revealed no changes in fibronectin deposition after the addition of polarised macrophages. This finding confirms that the observed increase in collagen deposition is not due to a generalised upregulation of extracellular matrix components (Supplementary Figure 2A). To further explore whether this fibril enhancing effect requires cell-cell contact, or through soluble factors secreted by macrophages, CM from polarised THP-1 macrophages was added to fibroblast cultures (Figure 3C). Of note, only CM from M2-polarised macrophages significantly enhanced collagen deposition in fibroblasts compared to lung fibroblast cultures without CM (Figure 3D).

**Figure 3.**
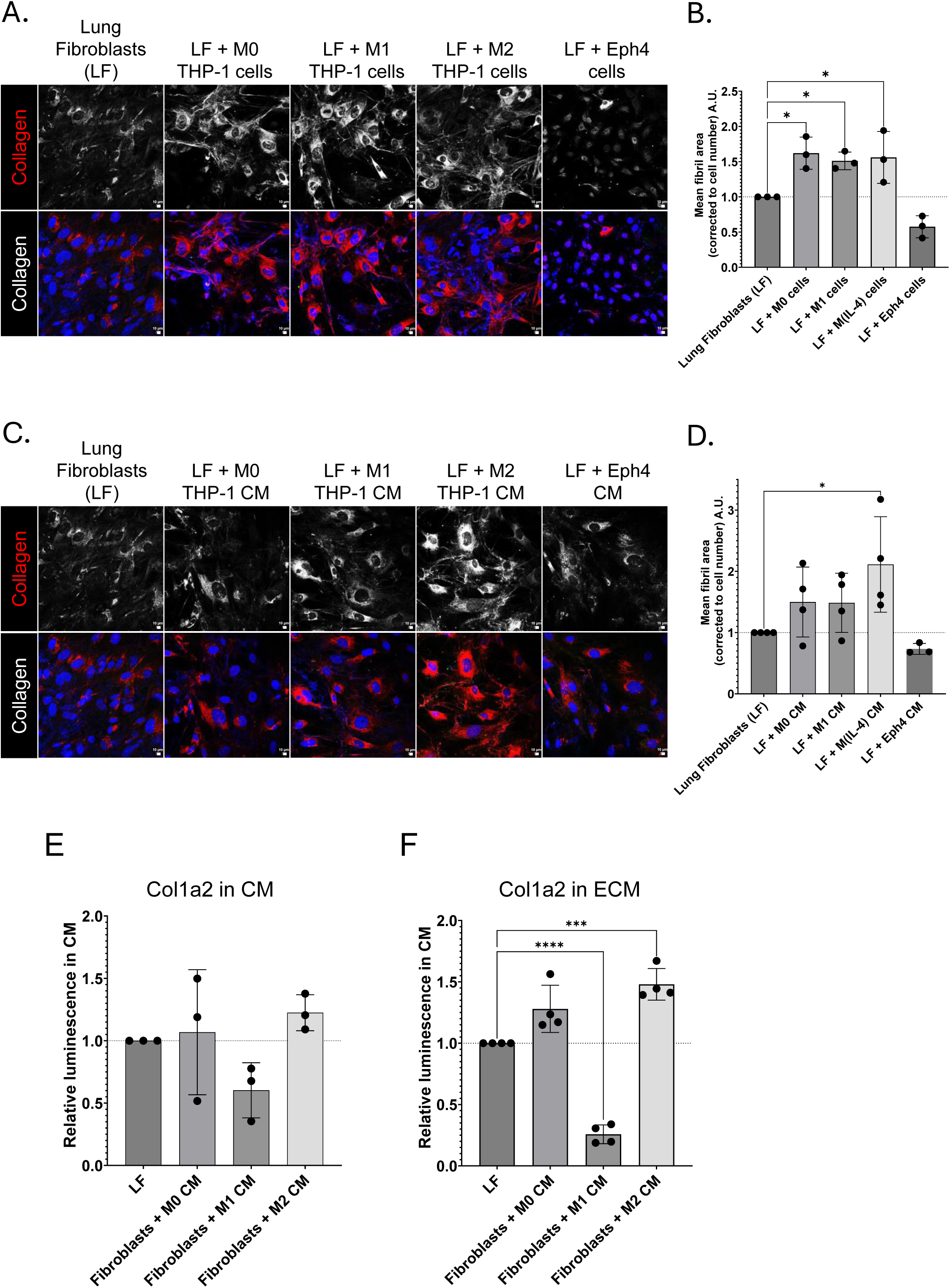
Quantification of collagen fibrils deposited by lung fibroblasts with or without THP-1 cells. (A) IF images of LF and THP-1 macrophage co-cultures. Top panels: collagen channel; bottom panels: pseudo-coloured collagen (red) and DAPI-stained nuclei (blue). Scale bar = 10 μm. Representative of N= 3. (B) Quantification of collagen fibril deposition in fibroblast cultures ± THP-1 macrophages. N = 3 biological repeats, *p<0.05, one-way ANOVA, error bars indicate standard deviation. (C) IF images of fibroblasts with THP-1 macrophage-conditioned medium (CM). Layout similar to Figure 3A. Scale bar = 10 μm. Representative of N= 4. (D) Quantification of collagen fibril area ± macrophage CM. N = 4 biological repeats for all bar LF + Eph4 CM, which is N = 3. *p<0.05, one-way ANOVA, error bars indicate standard deviation. (E) Primary murine HiBiT-Col1a2 fibroblasts were treated with CM from polarised primary AMs for 4 days before replacing with fresh media. Luminescence was measured in the media of these cultures after 48 hours. Luminescence is proportional to amount of Col1a2 in media. There was no significant difference in HiBiT luminescence readings between LF only and LF cultures treated with M0, M1 or M2 CM . N = 3 biological repeats, one-way ANOVA, error bars indicate standard deviation. (F) Luminescence readings were also taken in the culture wells of LF samples to assess luminescence and therefore Col1a2 deposition into the matrix. M2 CM significantly increased, while M1 CM significantly decreased the deposition of fibroblast-derived tagged Col1a2 into the ECM. N = 4 biological repeats, *** p<0.001, ****p<0.0001 one-way ANOVA, error bars indicate standard deviation.

Previously it was reported that macrophages can deposit collagen in mouse cardiac wound sites^4^. Here, we cultured M0, M1 and M2 THP-1 macrophages at different cell numbers for 4 days in the presence of ascorbic acid. Only cultures with over 80% confluency deposited collagen fibrils (Supplementary Figure 2B), indicating that whilst macrophages can produce collagen fibrils, it is unlikely that they are generating fibrils at the cell number used in our co-culture systems. To directly assess fibroblast specific collagen deposition, HiBiT-Col1a2 lung fibroblasts were used. These fibroblasts contain a genetically modified *Col1a2* gene, containing a bioluminescence-tagged collagen alpha-2(I) chain (Supplementary Figure 2C). To confirm the HiBiT-tag was incorporated into collagen fibrils, HiBiT-Col1a2 fibroblasts were imaged and stained using both antibodies against HiBiT and collagen. IF imaging showed co-localisation between HiBiT and collagen-I signals, indicating that the HiBiT-collagen is incorporated into fibrils as normal collagen (Supplementary Figure 2D). HiBiT-Col1a2 fibroblasts were then cultured with macrophage CM for four days. Collagen deposition was quantified by measuring relative luminescence of two distinct sampling: 1) collagen released into the media (CM, i.e. secreted protomeric collagen) and 2) collagen deposited into the ECM (i.e. collagen fibrils) (Figure 3E, F). Luminescence readings of fibroblast CM showed no significant changes in collagen release when cultured with macrophages (Figure 3E). However, luminescence readings of collagen deposited into the ECM revealed that M2 CM significantly increased collagen luminescence readings, indicating enhanced collagen deposition (Figure 3F). Interestingly, M1 CM resulted in a significant reduction in ECM collagen deposition.

To determine if the increased collagen deposition was due to elevated expression of collagen genes and thus protein production in fibroblasts, *Col1a1* gene expression was quantified by qPCR following the addition of polarised THP-1 macrophages and CM (Supplementary Figure 2E, F). Neither macrophage cells nor CM influenced *Col1a1* expression in fibroblasts, suggesting that the increase in collagen deposition is not through transcriptional control.

### Collagen deposition in fibroblasts is reliant on a functioning circadian clock

We have previously identified that collagen secretion and deposition in fibroblasts is dependent on circadian rhythms^5^. Here, to assess the impact of circadian dynamics on collagen fibril deposition, lung fibroblast cultures were analysed by IF over a 24-hour period at 4-hour intervals. Collagen deposition was found to exhibit rhythmicity in both fibroblast monocultures and co-cultures with THP-1 macrophages, whereby co-cultures had a larger collagen fibril area at the peak of fibril numbers (Figures 4A and 4B, respectively).

**Figure 4.**
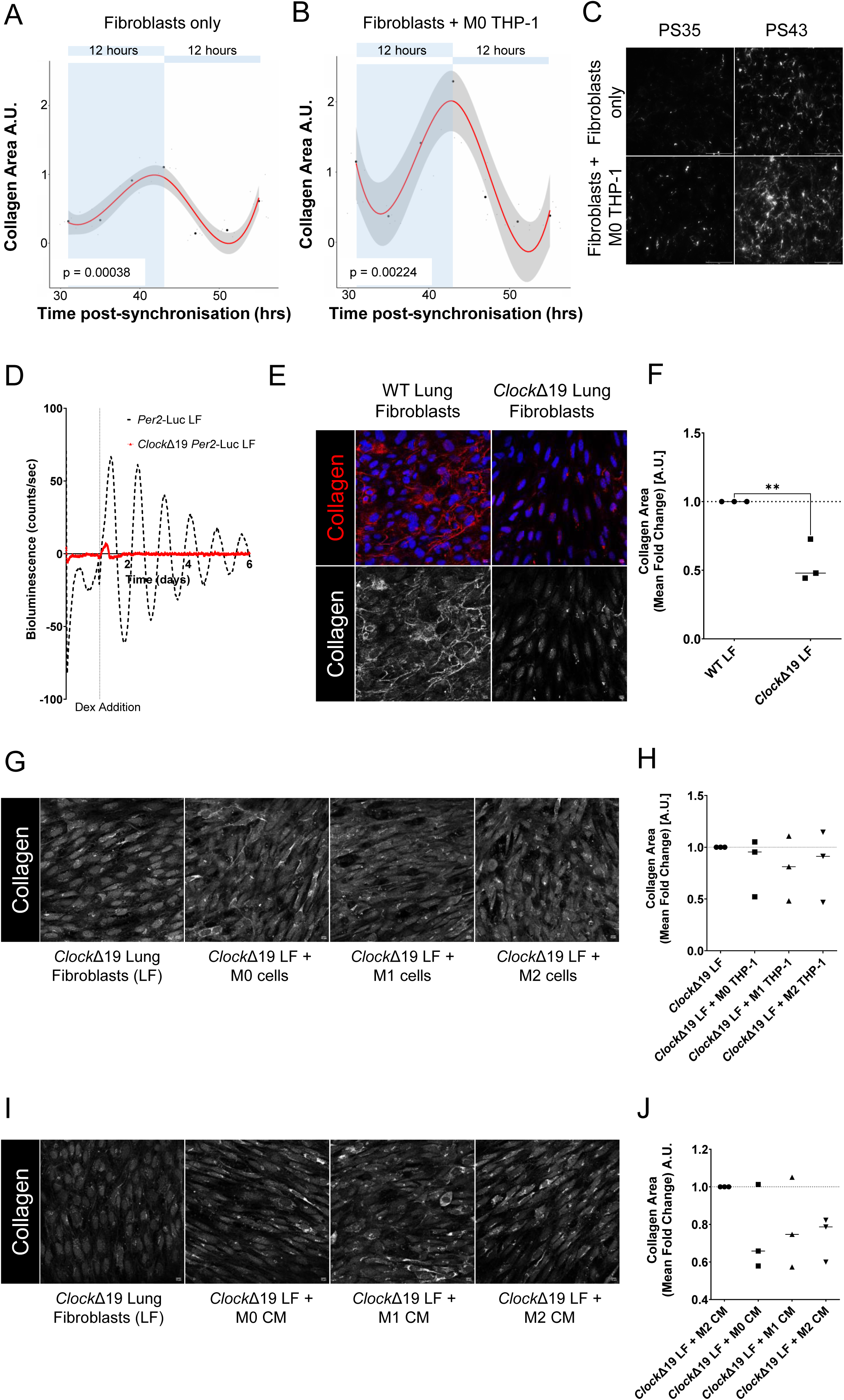
The Relationship Between Circadian Dynamics in Fibroblasts and Collagen Deposition. (A) Fibroblasts monoculture deposited collagen rhythmically over cultured with or without macrophages deposited collagen rhythmically over 24 hours after synchronisation with dexamethasone. Circadian analysis to determine rhythmicity was conducted using cosinor2 package in R to fit a cosinor model and assess rhythmicity. Representative of N= 3. ***p< 0.001, **p<0.01, p-value calculated using cosinor.detect() function on R. (B) Fibroblast co-cultured with macrophages deposited collagen rhythmically over 24 hours after synchronisation with dexamethasone. Circadian analysis to determine rhythmicity was conducted using cosinor2 package in R to fit a cosinor model and assess rhythmicity. Representative of N= 3. ***p< 0.001, **p<0.01, p-value calculated using cosinor.detect() function on R. (C) IF images of collagen fibrils at 35- and 43-hours post-synchronisation (PS) in lung fibroblast monocultures and co-cultures with M0 THP-1 macrophages. Representative of N= 3. (D) Bioluminescence readings indicated dexamethasone initiated a rhythm in *Per2*-Luc WT fibroblasts but not in *Clock*Δ19 fibroblasts. Representative of N= 3. (E) IF staining of collagen in *Clock*Δ19 and WT fibroblast cultures showed collagen (red) and nuclei (blue) in pseudo-color images (top) and grayscale collagen-only images. Scale bar = 10 μm. Representative of N= 3. (F) Quantification of collagen deposition in *Clock*Δ19 lung fibroblasts alone and WT lung fibroblasts showed that *Clock*Δ19 fibroblasts deposit significantly less collagen than WT lung fibroblasts. N = 3 biological repeats. (G) Representative IF images show *Clock*Δ19 fibroblasts alone and co-cultured with polarised macrophages. Representative of N= 3. (H) Polarised macrophages did not increase collagen deposition in *Clock*Δ19 fibroblasts, with similar collagen levels observed in monocultures and co-cultures. Scale bar = 10 μm. N = 3 biological repeats, *p<0.05, one-way ANOVA. (I) Representative IF images show *Clock*Δ19 fibroblasts alone and co-cultured with polarised macrophages. Representative of N= 3. (J) CM from polarised macrophages did not increase collagen deposition in *Clock*Δ19 fibroblasts . N = 3 biological repeats, *p<0.05, one-way ANOVA.

To further explore the role of the circadian clock in this process, fibroblasts with a truncated *Clock* gene and thus impaired circadian rhythm (*Clock*Δ19) were evaluated. Bioluminescence recordings showed negligible rhythmic activity apart from an initial peak following dexamethasone synchronisation (Figure 4C). Addition of macrophages did not rescue the rhythm in these *Per2*-Luc *Clock*Δ19 fibroblasts (Supplementary Figure 3).

Collagen deposition in *Clock*Δ19 fibroblasts was compared to WT fibroblasts. IF analysis revealed significantly reduced collagen deposition in *Clock*Δ19 lung fibroblast cultures (Figures 4D, E), suggesting a critical role for an intact circadian clock in regulating collagen synthesis and ECM dynamics in lung fibroblasts. To determine whether macrophages could compensate for the impaired circadian regulation of collagen deposition in *Clock*Δ19 fibroblasts, these cells were co-cultured with polarised THP-1 macrophages or treated with macrophage CM. IF images and quantification demonstrated that co-cultures with M0, M1, or M2 macrophages did not rescue collagen deposition in *Clock*Δ19 fibroblasts (Figures 4F, G). Similarly, the addition of M0, M1, or M2 macrophage CM to *Clock*Δ19 fibroblasts failed to increase collagen deposition (Figures 4H and 4I). Together, these findings highlight a reliance of fibroblasts on an intact circadian clock for ECM remodelling.

### PDGFA is highly upregulated in M2 conditioned media□□

As M2 CM induces the most robust circadian rhythm and enhances collagen deposition in fibroblasts, we aimed to identify the factor(s) responsible for this change. CM from M0, M1, and M2 THP-1 macrophages were analysed by MS to identify proteins significantly upregulated in M2 CM compared to M0 and M1 CM. Proteins identified through MS were clustered using k-means clustering, resulting in two distinct clusters. Proteins were considered significantly differentially expressed if they exhibited a fold-change (FC) ≥ 2 and a p-value ≤ 0.05.

Cluster 1 comprised of proteins upregulated in M1 CM compared to M0 and M2 CM (Figure 5A). Established known M1-expressed proteins in this cluster include CXCL8 and CXCL10^29^, which were among the most differentially expressed genes compared to M2 CM, confirming the polarisation status of the macrophages. Cluster 2 contained proteins upregulated in M2 CM, with M2 macrophages showing greater proteomic similarity to M0 than to M1 (Figure 5B). Amongst the proteins elevated in M2 CM, PDGFA emerged as the most highly differentially expressed, displaying strongly positive log2 fold-change values when compared to both M0 and M1 CM.

**Figure 5.**
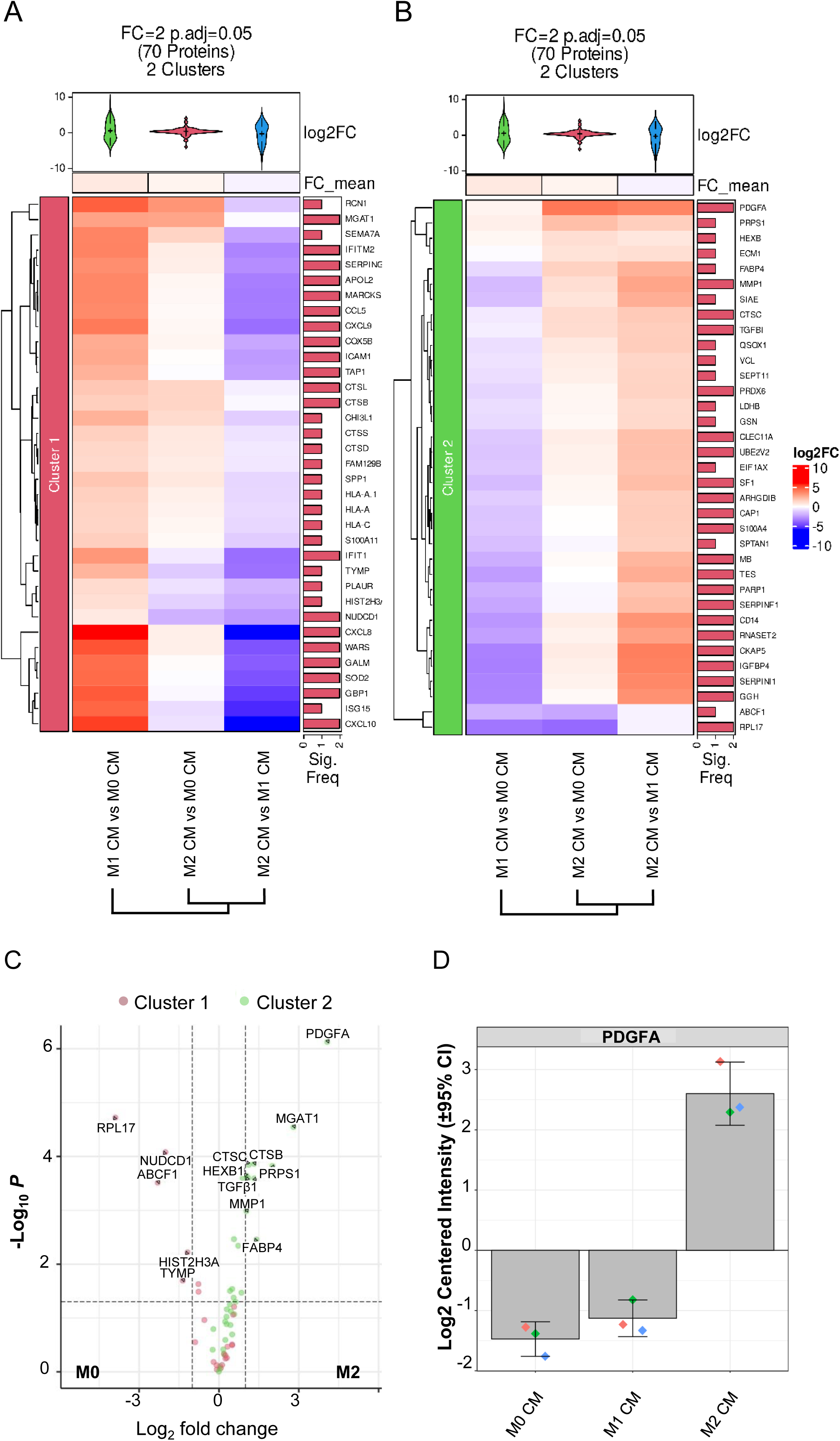
Comparative Proteomic Analysis of Differentially Expressed Proteins in M0, M1, and M2 AM CM. (A) Heatmap analysis of differentially expressed proteins in M0, M1, and M2 AM CM, based on MS data processed with the Manchester Proteome Profiler. Fold change values (≥ 2, p ≤ 0.05) were clustered using k-means. Cluster 1 highlights proteins upregulated in M1 CM, while cluster 2 shows proteins upregulated in M2 CM. N = 3 biological repeats. (B) Cluster 2 includes PDGFA, which is highly expressed in M2 CM but not M0 or M1 CM. N = 3 biological repeats. (C) Volcano plot of proteins in M2 CM vs. M0 CM. Proteins significantly upregulated in M0 CM appear on the left, while those upregulated in M2 CM, including PDGFA as the most prominent, appear on the right. N = 3 biological repeats. (D) Barplot showing positive log2 centered intensity in M2 CM samples only. N = 3 biological repeats.

To visualise relative protein expression, volcano plots were generated using the Manchester Proteome Profiler. These plots applied the same FC and p-value thresholds, with dashed lines marking the cut-offs. Proteins differentially expressed between M2, and M0 CM are highlighted in Figure 5C, where green dots indicate proteins elevated in M2 CM and pink dots represent those higher in M0 CM. PDGFA appeared prominently in the top-right quadrant, signifying significant upregulation in M2 CM. Additional analysis comparing M2 to M1 CM, and M1 to M0 CM, were performed (Supplementary Figure 4A, B).

The abundance of PDGFA in M2 CM was further emphasised when the data is presented as log2-centered intensity (Figure 5D), where positivity was exclusively observed in M2 CM samples. These findings highlight PDGFA as a key M2 macrophage-derived factor that may be driving the observed effects on fibroblast circadian dynamics and collagen deposition.

As PDGFA has relatively high thermal stability^30^, CM from THP-1 macrophages was also subject to heat-treatment. The media was heated to 95°C for 30 minutes, cooled to room temperature for 30 minutes, and then added to *Per2*-Luc lung fibroblasts. Heat-treated CM induced a circadian rhythm in the fibroblasts similar to that of untreated CM. Quantification of circadian amplitude revealed no significant differences between heat-treated and untreated CM (Supplementary Figure 4C, D), confirming heat-stability in the rhythm-inducing soluble factor(s) and providing further evidence that PDFA is likely involved in driving circadian changes in fibroblast cultures.

### Macrophage-derived PDGFA Enhances Collagen Deposition and Increases Circadian Rhythm Amplitude in Fibroblasts□

PDGFA was investigated as a candidate protein responsible for enhancing collagen deposition and circadian rhythm amplitude in fibroblasts. We first approached this using recombinant PDGFA at a concentration of 17.5 ng/mL, consistent with what has been measured in whole blood serum^31^. Addition of recombinant PDGFA to *Per2*-Luc fibroblasts significantly increased circadian amplitude, as shown in Figures 6A and 6B. We then performed CRISPR on THP-1 cells to knock-out PDGFA expression (PDGFAko). The knock-out was confirmed by sequencing and western blot (Supplementary Figure 5A, B). Furthermore, unlike WT THP-1 macrophages, PDGFAko THP-1 did not induce a sustained rhythm in *Per2*-Luc fibroblasts (Figure 6C, D).

**Figure 6.**
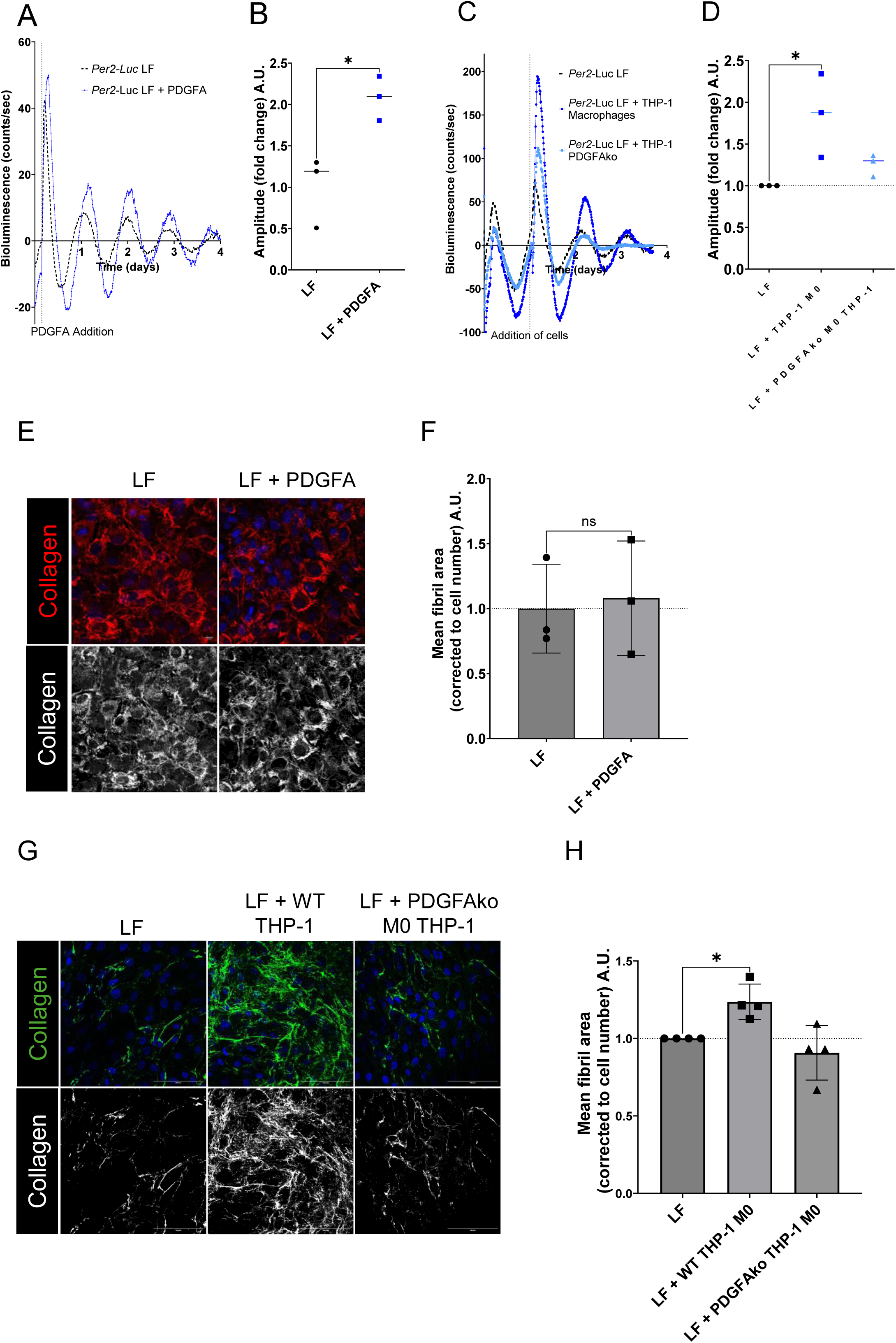
Evaluation of PDGFA’s capability to enhance circadian amplitude and collagen deposition in fibroblasts. (A) Recombinant PDGFA induced a robust circadian rhythm in *Per2*-Luc lung fibroblasts. Representative of N= 3. (B) Amplitude fold change analysis demonstrated that PDGFA significantly enhances circadian amplitude. N = 3 biological repeats, **p<0.01, one-way ANOVA, error bars indicate standard deviation. (C) THP-1 PDGFAko failed to induce a sustained and robust circadian rhythm in *Per2*-Luc lung fibroblasts. Representative of N= 3. (D) Amplitude fold change analysis showed that WT THP-1 M0 significantly increased circadian amplitude in fibroblasts, while PDGFAko THP-1 M0 did not. N = 3 biological repeats, **p<0.05, un-paired two-tailed t-test, error bars indicate standard deviation. (E) IF images of lung fibroblasts in with recombinant PDGFA. Top panels show composite images of DAPI-stained nuclei (blue) and collagen (red), while bottom panels show the collagen channel only. Scale bar = 10 µm. (F) Quantification revealed there was no statistical difference between collagen deposition in fibroblast cultures following incubation with PDGFA. N = 3 biological repeats, error bars indicate standard deviation. (G) IF images of lung fibroblasts in monoculture and co-culture with either PDGFA KO THP-1 or WT THP-1 macrophages. Top panels show composite images of DAPI-stained nuclei (blue) and collagen (green), while bottom panels show the collagen channel only. Representative of N=4. Scale bar = 10 µm. (H) Quantification revealed that WT THP-1 macrophages significantly increased collagen deposition in fibroblasts, whereas PDGFA KO THP-1 macrophages did not. N = 3 biological repeats, *p<0.05, one-way ANOVA, error bars indicate standard deviation.

To assess whether recombinant PDGFA alone could stimulate collagen deposition in fibroblast cultures, imaging studies were conducted. As shown in Figure 6E, no significant difference in collagen deposition was observed when fibroblasts were treated with recombinant PDGFA compared to untreated controls. Quantitative analysis of collagen area further confirmed that PDGFA alone did not influence collagen deposition in fibroblasts (Figure 6F). IF imaging was also carried out to compare the effects of PDGFAko and WT THP-1 macrophages on collagen deposition (Figure 6G). Consistent with previous findings, fibroblasts co-cultured with WT THP1 deposited a significantly increased amount of collagen fibrils when compared to fibroblast monoculture (middle panel vs. left panel), whereas fibroblasts co-cultured with PDGFAko THP-1 macrophages have similar levels of collagen fibrils compared to monoculture (right vs. left panel), as confirmed by quantification (Figure 6H). This indicates that macrophage-derived PDGFA drives increase of collagen fibrillogenesis.

Taken together, these results identify PDGFA as a key macrophage-derived factor that enhances collagen deposition and circadian rhythm amplitude in fibroblasts, highlighting it as a mediator of macrophage-fibroblast crosstalk.□

## Discussion

In this study, we provided evidence that polarised macrophages, as well as soluble factors secreted by M2-polarised macrophages, enhanced collagen deposition in fibroblasts via a clock-mediated mechanism. These results highlight the importance of effective crosstalk between macrophages and fibroblasts, as well as demonstrating the necessity of a functioning clock, in order to maintain a functional ECM. These data therefore have implications for tissue remodelling and repair processes. It is important to note that, in general, “M2-polarised macrophages” refers to multiple different subtypes with certain features in common, such as being anti-inflammatory, but the specific role and characteristics depend on the cell’s exposure to different cytokines and growth factors etc. In this study, “M2-polarised macrophages” refers only to naïve M0 macrophages polarised with either IL-4c (*in vivo*) or IL-4 and IL-13 (*in vitro*).

While M0, M1 and M2 THP-1 macrophages all increased collagen deposition and circadian amplitude in co-cultures with fibroblasts, only CM from M2-polarised macrophages had the same effect. This is consistent with M2 macrophages’ phenotype, which involves ECM remodelling and pro-wound healing characteristics^32,33^. These findings therefore suggest that macrophages regulate fibroblast function via both paracrine and juxtacrine signalling pathways. However, M2-polarised macrophages have an increased capacity to regulate collagen deposition in fibroblasts, via elevated secretion of circadian-modulating soluble factors – a mechanism that would allow M2 macrophages to drive ECM remodelling.

We have previously identified that fibroblasts utilise endocytic recycling as a mechanism for collagen fibrillogenesis^34^, and prior research has identified that macrophages can directly deposit collagen during heart repair^4^. These observations suggest that the source of collagen in a complex environment may not be strictly from fibroblast-like cells. Here we observed that macrophages, regardless of polarisation status, can produce collagen fibrils, although morphologically different from fibroblast-derived fibrils and only observed at high cell density. As the number of macrophages in our co-culture systems was much lower than fibroblasts, and macrophages are terminally differentiated cells that do not replicate *in vitro*, it is highly likely that the collagen fibrils are produced by fibroblasts only, although we cannot rule out that macrophage-derived collagen may have been taken up by fibroblasts and utilised for fibrillogenesis. This could be addressed in the future by utilising an endogenous collagen-specific HiBiT tag in macrophages and co-culturing with fibroblasts. The bioluminescence signal could then be used to determine the amount of collagen within the ECM derived from macrophages.

*In vitro* polarised M0 and M2 AMs both enhanced circadian amplitude in fibroblasts, whereas only *in vivo* polarised M2 macrophages induced a similar effect. These differences may stem from the inherent variability of primary macrophages, influenced by their origin and microenvironment. Nevertheless, M2 macrophages consistently demonstrated a significant ability to induce circadian rhythms in fibroblasts, highlighting their heightened capacity for intercellular communication and modulation of fibroblast circadian dynamics. Interestingly, we also observed that M2-secreted factors in particular significantly induced fibroblast circadian- and collagen-response. Mass spectrometry analysis on the secretome of M0, M1, and M2 macrophages identified over 1000 proteins, which was further filtered for decoys and known contaminants, and analysed in a pairwise manner (M0 vs M2, M1 vs M2) to identify 35 of the most significantly differentially expressed (i.e. most up-regulated and most down-regulated) proteins for each comparison. PDGFA was identified as the top upregulated protein in M2 CM compared to M0 CM and M1 CM, suggestive that it is a key macrophage-secreted factor that drives increased collagen deposition, through a clock-dependent mechanism. Additional support for this hypothesis comes from the observation that heat treatment of M2 CM did not reduce its ability to enhance circadian amplitude in fibroblasts, consistent with the known thermal stability of PDGFA^30^. In our studies, recombinant PDGFA significantly enhanced circadian amplitude in fibroblasts. Furthermore, while WT THP-1 macrophages increased circadian amplitude in fibroblasts, THP-1 macrophages lacking PDGFA (PDGFAko) failed to produce the same effect. Hence, PDGFA was demonstrated to be essential for enhancing circadian amplitudes in co-culture systems.

Although PDGFA was found to be highly upregulated exclusively in CM from M2-polarised macrophages, unpolarised PDGFAko M0 THP-1 macrophages were unable to enhance collagen deposition in fibroblasts, unlike their WT counterparts. This suggests that even low levels of PDGFA may be essential for facilitating effective cell-cell interactions between macrophages of all phenotypes and fibroblasts. Supporting this hypothesis, our data demonstrate that both M0 and M1 macrophages require direct cell contact to increase collagen fibril deposition in fibroblasts. This aligns with a previous study showing that PDGFA functions as a potent chemoattractant for fibroblasts^35^. In contrast, CM from M2 macrophages alone is sufficient to promote collagen fibril deposition. Notably, however, recombinant PDGFA alone, added at a concentration consistent with levels detected in whole blood serum, was unable to increase collagen deposition in fibroblasts. This is an unexpected finding, as previous data has shown that mice with a PDGFRα KO exhibited significantly reduced collagen deposition, indicating that PDGFA signalling plays a crucial role in collagen regulation^36^. The disparity in findings may be indicative of the necessity for an elevated concentration of PDGFA in *in vitro* cultures.

Taken together, this suggests that PDGFA synchronises the circadian clock in fibroblasts, but on its own is not sufficient to increase collagen deposition in fibroblasts, suggestive of a requirement for other macrophage-derived factors working in tandem. Regardless, these findings point to a crucial role for PDGFA in synchronising fibroblast circadian rhythms, which may subsequently influence the temporal regulation of clock-controlled genes involved in the collagen secretory pathway. Although beyond the scope of the present study, future studies should aim to identify additional components of M2 CM that work with PDGFA to augment fibroblast collagen deposition. These efforts could involve proteomic and transcriptomic analyses to characterise the secretome of M2 macrophages, with or without PDGFA, comprehensively.

An alternative, but not mutually exclusive, explanation is that PDGFA-mediated circadian synchronisation enhances the responsiveness of fibroblasts to other macrophage-derived factors. The circadian clock regulates the expression of genes involved in the collagen secretory pathway^5^. By amplifying the circadian amplitude, PDGFA may prime fibroblasts for optimal temporal coordination of these clock-controlled genes, thereby facilitating increased collagen deposition when additional signals are present.

Collectively, our findings suggest that M2 macrophages secrete multiple factors that drive fibroblast circadian synchronisation and subsequent collagen deposition. While PDGFA is a key mediator, the presence of additional soluble factors is likely critical to the observed effects. Further studies are warranted to identify and characterise these factors and to elucidate their individual and collective contributions to circadian regulation and ECM remodelling. Insights from these studies could guide the development of therapeutic strategies aimed at modulating macrophage-fibroblast interactions to optimise tissue repair mechanisms.

## Methods

### Mouse Experimentation

The care and use of all mice was carried out in accordance with the UK Home Office regulations, UK Animals (Scientific Procedures) Act of 1986 under the Home Office Licence number, S1-P57F83603 and P44492AC9.

All lung fibroblasts (LFs) and alveolar macrophages (AMs) were isolated from C57BL/6J mice. *Per2*-Luc lung fibroblasts were derived from C57BL/6J mice bred on a background with PER2::Luciferase reporter^14^. *Clock*Δ*19* mice (C57BL/6 on a *Per2*:Luciferase reporter background) were as described previously^15^.

*In vivo* AM polarisation was carried out by administering cytokines into the nostrils of 12-week-old mice, under the Home Office Licence number P44492AC9. LPS was used to induce M1 polarisation (0.15 µg in 30 µL PBS). Interleukin-4c (IL-4c) was utilised to induce M2 polarisation in alveolar macrophages. The IL-4c was prepared by combining recombinant IL-4 (PeproTech, 214-14) with an anti-IL-4 antibody (BD Biosciences, 554387) in a 1:5 ratio of IL-4 to antibody. Specifically, 0.05 µg of IL-4c was diluted in 30 µL of phosphate-buffered saline (PBS) for administration. Mice were given two doses of either PBS, IL-4c or LPS on days 0 and 2 under isoflurane-induced anaesthesia. Mice were then culled by rising CO_2_ on day 4.

### Primary cell isolation

Primary lung fibroblasts were isolated from mice aged between 2 and 3 months. Tissue was digested in 1000 U/ml bacterial collagenase type 4 (Worthington Biochemical Corporation) in trypsin (2.5 g/L), as previously described^16^.

AMs were isolated from 3-month-old mice by broncho-alveolar lavage (BAL), as previously described^17^.

### Cell culture and *in vitro* cell differentiation

*In vitro* THP-1 monocytes were differentiated into macrophages by 48 hours incubation with 100 nM phorbol 12-myristate 13-acetate (PMA, Sigma Aldrich, P8139). Macrophages were polarised to M1 macrophages by incubation with 20 ng/ml of IFN-γ (Miltenyi Biotec, 130-096-481) and 10 ng/ml of LPS (Sigma-Aldrich, 297-473-0) for 24 hours. Macrophages were polarised to M2 phenotype by incubation with 20 ng/ml of IL-4 (Miltenyi Biotec, 130-093-920) and 20 ng/ml of IL-13 (Miltenyi Biotec, 130-112-409) for 24 hours.

All macrophages were maintained in culture in Roswell Park Memorial Institute (RPMI 1640, Sigma-Aldrich, R8758) culture medium containing 2 mM L-glutamine and 1.5 g/L sodium bicarbonate. The media was supplemented with 10% heat inactivated fetal bovine serum (FBS), 1% penicillin and streptomycin antibiotics (Sigma-Aldrich, P4333), 10 mM Hepes (Gibco, 15630-080) and 1 mM sodium pyruvate (Gibco, 11360-039).

All fibroblasts were maintained in Dulbecco’s Modified Eagle’s Medium F-12 (DMEM F-12, Gibco, 11320033) culture medium containing 2 mM L-glutamine. The media was supplemented with 10% heat inactivated FBS and 1% antibiotics. Cells were maintained at 37°C in a 5% CO_2_ humidified incubator.

Eph4 cells were maintained in Dulbecco’s Modified Eagle’s Medium (DMEM) with high glucose culture medium (Sigma-Aldrich, D5796), containing 4500 mg/L glucose, L-glutamine, and sodium bicarbonate, supplemented with 10% heat inactivated FBS and 1% antibiotics.

Unless otherwise stated, during experimentation all cells were cultured in the presence of 200 µM L-ascorbic acid.

### Preparation and collection of Conditioned Media (CM)

Cells were cultured for at least 5 days in serum-free media. CM was taken each day and passed through a 0.45 µM filter to remove cells and cell debris. The CM was then frozen until use.

### Generation of PER2-Luc2 THP-1 cell line

PER2-Luc2 lentivirus was produced using HEK293T cells and transduced into THP-1 cells. Briefly, the PER2-Luc2 encoding lentivirus was produced by adding 1 mL Opti-MEM (Thermo Fisher Scientific, 31985062) to an Eppendorf with 2.5 µg psPAX2 packaging vector (Addgene, 12260), 2.5 µg pMD2.G envelope (Addgene,12259) and 5 µg per2-bre-luc2-ef1a-GFP transfer plasmid, which has been previously generated^18^. This mixture was incubated at room temperature for 15 minutes. HEK293T cells were transfected with the lentivirus and incubated for at least 48 hours before the virus was harvested. The viral supernatant was centrifuged at 500 x g, 4°C, for 5 minutes to pellet any remaining cells and then passed through a 0.45 µm polyethersulfone membrane filter. Virus transduction into THP-1 cells was achieved by incubating cells in a 6-well plate with 1 mL of the virus suspension and 8 µg/µL polybrene (Sigma Aldrich, TR-1003). The cells were incubated with the virus suspension for 48 hours. The plasmid also contained a GFP tag, so fluorescence was observed using fluorescent microscopes to ensure that transduction of the plasmid was successful.

### Generation of endogenously-tagged HiBiT-collagen lung fibroblasts

HiBiT-Col1a2 CRISPR-edited lung fibroblasts were genetically modified to endogenously express luminescently-tagged Col1a2 with a HiBiT tag at the end of the N-propeptide^19^. This strategy for editing these cells has been shown to preserve collagen function while serving as an effective quantitative reporter for collagen^20^.

### Generation of PDGFA knockout (KO) in THP-1 cell line

Cells were treated with CRISPR-Cas9 to delete the PDGFA gene. Two sgRNAs targeting PDGFA were designed, with sequences: cgggcacatgcttagtggca (g1) and catggaccccgtgagctctc (g2). The sgRNAs were introduced into the genome of THP-1 cells using CRISPR-Cas9. sgRNA and tracrRNA were combined in a 1:1 ratio before heating at 95°C for 5 minutes. The mixture was cooled to room temperature before centrifuging at 10,000 x g for 30 seconds. Cas9 was then added to the mixture at a 1:2 ratio and mixed. This was left for 10 minutes at room temperature to allow for annealing. The RNP complex was then delivered into THP-1 cells via electroporation using Neon NxT electroporation system (Thermo Fisher Scientific).

### Circadian bioluminescence recording

The circadian rhythm of cells isolated from the *Per2*-Luc mouse and macrophages containing the PER2-Luc2 plasmid were measured using LumiCycle apparatus (Actimetrics), which provided real-time quantitative bioluminescence recordings. Traces were typically recorded for a minimum of 4 days. The LumiCycle analysis software (Actimetrics) performed baseline subtraction of recorded bioluminescence values using a 24-hour moving average. 100 nM dexamethasone (dex) (Scientific Laboratory Supplies, D1756) was added to synchronise the cells’ circadian rhythm. It was added to cell cultures in the LumiCycle following 24 hours of recording (unless otherwise shown in figure) to establish how the circadian rhythm of a synchronised population should look like. Amplitude was calculated using BioDare circadian analysis software.

### HiBiT::Col1a2 luminescence readings

Primary murine lung fibroblasts expressing HiBiT::Col1a2 were cultured in 96-well plates (10,000 cells per well) for 48 hours before addition of macrophage CM containing a total of 25 µg of protein. Fibroblasts were incubated with the CM for 4 days. CM was then removed, and cells were washed extensively with PBS before adding fresh media to determine collagen secretion. HiBiT::Col1a2 luminescence readings commenced after 48 hours. Nano-Glo® HiBiT Extracellular Detection Kit (Promega, N2420) and a microplate reader (SYNERGY neo2, BioTek) were used to measure HiBiT-tagged Col1a2 in both the media from lung fibroblast culture wells and the wells themselves, thus capturing secreted and deposited Col1a2, respectively.

### Mass spectrometry proteomics analysis of CM

25 µg of protein per sample was prepared for Mass Spectrometry (MS)-based proteomics analysis using S-Trap columns (Protifi, C02-micro-10) where sequencing grade modified trypsin (Promega, V5111) was used for digestion following a previously published protocol^21^. Porous Oligo R3 beads (Thermo Fisher Scientific, 1133907) were used to desalt peptides. Peptides were then dried in a vacuum centrifuge and resuspended in 10 µL of a 5% acetonitrile, 0.1% formic acid solution for MS analysis.

Liquid chromatography separation was performed on a Thermo RSLC system consisting of a NCP3200RS nano pump, WPS3000TPS autosampler and TCC3000RS column oven configured with buffer A as 0.1% formic acid in water and buffer B as 0.1% formic acid in acetonitrile. An injection volume of 2 ul was loaded into the end of a 5 ul loop and reversed flushed on to the analytical column (Waters nanoEase M/Z Peptide CSH C18 Column, 130Å, 1.7 µm, 75 µm X 250 mm) kept at 35 °C at a flow rate of 300 nl/min for 8 min with an initial pulse of 500 nl/min for 0.3 min to rapidly re-pressurise the column. The injection valve was set to load before a separation consisting of a multistage gradient of 1% B to 6% B over 2 minutes, 6% B to 18% B over 44 minutes, 18% B to 29% B over 7 minutes and 29% B to 65% B over 1 minute before washing for 4 minutes at 65% B and dropping down to 2% B in 1 minute. The complete method time was 75 minutes.

The analytical column was connected to a Thermo Orbitrap Fusion Lumos Tribrid mass spectrometry system via a Thermo nanospray Flex Ion source via a 20 um ID fused silica capillary. The capillary was connected to a stainless steel emitter with an outer diameter of 150um and an inner diameter of 30 um (Thermo Scientific, ES542) via a butt-to-butt connection in a steel union. The nanospray voltage was set at 2000 V and the ion transfer tube temperature set to 275 °C.

Data was acquired using ion trap MS/MS in a data dependent manner using a fixed cycle time of 2 sec, an expected peak width of 15 sec and a default charge state of 2. Full MS data was acquired in positive mode over a scan range of 375 to 1500 Th, with a resolution of 120,000, a normalised AGC target of 100%, in automatic injection time mode for a single microscan. Fragmentation data was obtained from signals with a charge state of +2 to +4 and an intensity over 5,000 and they were dynamically excluded from further analysis for a period of 15 sec after a single acquisition within a 10ppm window. Fragmentation spectra were acquired in ion trap mode, with a normalised collision energy of 30%, a normalised AGC target of 100%, first mass of 110 Th and a max fill time of 35 mS for a single microscan. All data was collected in centroid mode.

Mass spectroscopy raw data were processed using MaxQuant version 2.4.10.0 for data-dependent acquisition and label-free quantification (LFQ), using the default settings unless stated otherwise. Database searches were conducted using the Andromeda search engine with human UniProt sequences (UP000005640, accessed November 13, 2023) as a reference and a contaminants database of common laboratory contaminants included with MaxQuant. Oxidation of methionine and protein N-terminal acetylation were specified as variable modifications and cysteine carbamidomethylation was specified as fixed modification. A minimum peptide length of seven was specified and the maximum peptide mass was limited to 4600 Da with a maximum of two missed cleavage sites. First search and main search peptide tolerance were set to 20 ppm and 5 ppm, respectively, and ITMS MS/MS match tolerance, de novo tolerance, and deisotoping tolerance were set to 0.5 Da, 0.5 Da, and 0.3 Da, respectively. Peptide spectrum matches and protein FDR was set to 1%, with “razor protein FDR” and “match between runs” selected. Mass spectrometry outputs (as LFQ intensity values) were further analysed using Perseus version 2.0.11. Data were initially filtered to remove contaminants and decoy sequences, log2 transformed, and then samples grouped into experimental groups. Data were further filtered to remove entries in which none of the groups had at least two numeric values. Remaining “NaN” values generated by the log2 transformation were replaced by imputation from theoretical normal distribution, with a width of 0.3 and a down shift of 1.8. Inter-group differences were assessed using one-way ANOVA followed by post-hoc Turkey’s HSD test. Results with False Discovery Rate (FDR) less than 0.05 were considered significant. Further analysis and figure creation were carried out using Manchester Proteome Profiler (Web-based analysis platform developed by Martin Humphries and Stuart Cain, University of Manchester, manuscript in preparation. Contact for access/information). R packages were used to process protein intensity data and cluster analysis was performed to allow data visualisation and identification of significantly enriched proteins. The DEP2 R package, which performs contaminant filtering, normalisation and imputation of missing values was used. Normalisation was performed using variance stabilising transformation (VSN), with missing values imputed using the Missing Not At Random method (MNAR). A p-value of 0.05 and a fold change of 2 were chosen and applied to determine a list of 35 most significantly regulated proteins in each pairwise comparison. Mass spectrometry data was deposited onto PRIDE with the identifier of PXD061123.

### RNA extraction, cDNA synthesis and quantitative PCR

To lyse cells for RNA extraction, cells were washed with PBS and snap frozen in liquid nitrogen with 2500μL TRIzol (Thermo Fisher Scientific, 15596026). RNA was isolated according to manufacturer’s instructions and RNA concentration was measured using NanoDrop One (Thermo Fisher Scientific). cDNA synthesis was performed using the TaqMan Reverse Transcription Kit (Thermo Fisher Scientific, N8080234). 500 ng – 1000 ng of RNA was used per sample for cDNA synthesis, where the same RNA amount was used in all samples within the same experiment.

The qPCR master mix was made using 2X SYBR Green PCR Master Mix (Thermo Fisher Scientific, 4309155), 10 µM primer mix (forward and reverse) and nuclease-free water before addition of cDNA. Mouse primer sequences (Sigma-Aldrich) were *18s* GTAACCCGTTGAACCCCATT and CCATCCAATCGGTAGTAGCG, *Cd64* GTCGGTGGGGAAGTGGTTAAAT and CCCCTCACACCATAAAGTGAC, *Col1a1* GCCTGCTTCGTGTAAACTCC and TTGGTTTTTGGTCACGTTCA, *Relm-*α CAAGGAACTTCTTGCCAATCCAG and CCAAGATCCACAGGCAAAGCCA. Human primer sequences were 18S GAGACTCTGGCATGCTAACTAG and GGACATCTAAGGGCATCACAG, CD64 AACTCTGCTCCTTTGGGTTCC and CTCTTGGAACACGCTGACCC, TGF-β1 CACGTGGAGCTGTACCAGAA GAACCCGTTGATGTCCACTT.

### Protein Extraction and Western Blotting

CM was generated as described above, and protein concentration determined by Bradford assay. 25 µg of protein was used for western blot analysis. Briefly, samples were separated using SDS-PAGE using 4-12% Bolt™ Bis-Tris gel (Thermo Fisher Scientific). This is followed by transfer to nitrocellulose membranes using Biorad TransBlot® Turbo^™^, blocked with 5% BSA in PBS for an hour, before incubation with PDGFA primary antibody (Merck, EMU041741) at 1:500. This was followed by incubation with secondary antibody (Invitrogen, A21058) at a concentration of 1:10,000. The bound antibody signals were then detected using Licor CFX system. Ponceau red stain was used to confirm equal loading of samples.

### Immunofluorescence, staining, microscopy and analysis

Cells plated on coverslips or 8-well μ-slides (Ibidi) were fixed with 100% methanol at -20°C and then permeabilized with 0.2% Triton-X in PBS. Primary antibodies used were as follows: rabbit pAb collagen (1:400, Gentaur OARA02579), mouse mAb FN1 (1:400, Sigma F6140), mouse HiBiT pAb (Promega; CS2006A01). Secondary antibodies conjugated to Alexa Fluor 488, Alexa Fluor 647, Cy3, and Cy5 were used (Thermo Scientific), and nuclei were counterstained with DAPI (Sigma). Coverslips were mounted using Fluoromount G (Southern Biotech).

Cells were imaged using an SP8 inverted confocal microscope (Leica) with a HC PL Apo 40x/1.30 oil objective. Settings were as followed: pinhole, 1□ Airy unit; scan speed, 400 Hz bidirectional; format, 1,024 × 1,024. Sequential collection of images was used to eliminate crosstalk between channels. Quantification of collagen fibrils was done using ImageJ software and achieved using the previously explained method^22^. Briefly, collagen fibrils were quantified by splitting the acquired image into its separate channels and adjusting the threshold value so that only collagen fibrils were present. Once set, the threshold value remained consistent in all images. The area of collagen fibrils was then measured and normalised to nuclei number. Nuclei were counted using CellProfiler software.

### Statistics and Reproducibility

Data are presented as mean ± SD unless otherwise specified in the figure legends. The sample size (N) represents the number of independent biological samples used in each experiment. Details regarding sample size, experimental repeats, and statistical analyses are provided in the figures, figure legends, or the Methods section. Data analysis was performed as described in the figure legends and was not conducted under blinded conditions.

Statistical significance was defined as p < 0.05. The significance levels are denoted as follows: *p < 0.05; **p < 0.01; ***p < 0.001; ****p < 0.0001; and n.s. (not significant). Actual p-values are reported where applicable. All analyses were conducted using GraphPad Prism 10 software.

## Author contributions

JC, TH and KEK conceived the project. JC, KEK, QJM, TH, AM supervised the experiments. KL, MFAC, SL, JC, MC performed experiments and interpreted data. JH performed *in vivo* polarization experiments. JK performed proteomics experiments. AM, QJM, SL provided reagents and data interpretation. All authors contributed to writing the manuscript.

## Acknowledgements

The proteomics was performed at the Biological Mass Spectrometry Facility with the assistance of Stacey Warwood, imaging was performed at the Bioimaging Facility, the PDGFA knock-out line was generated by the Genome Editing Unit; all facilities are in the Faculty of Biology, Medicine and Health (University of Manchester), and supported by funding from Wellcome, BBSRC, and the University of Manchester Strategic Fund. We thank the University of Manchester Biological Support Facility for expertise and assistance in animal welfare and husbandry.

## Funding

MRC Career Development Award MR/W016796/1 (JC)

Wellcome Trust Immunomatrix in Complex Disease (ICD) PhD studentship 218491/Z/19/Z (KL)

BBSRC strategic Longer and Larger Grants BB/T001984/1 (MFAC, QJM, KEK) Wellcome 226804/Z/22/Z (JC, QJM, GEU)

## References

1. Kadler, K.E. et al. Collagens at a glance. J. Cell. Sci. 120, 1955–1958 (2007).

2. Plikus, M. V. et al. Fibroblasts: Origins, definitions, and functions in health and disease. Cell 184, 3852–3872 (2021).

3. Wynn, T. A., Chawla, A. & Pollard, J. W. Macrophage biology in development, homeostasis and disease. Nature 496, 445–455 (2013).

4. Simões, F.C. et al. Macrophages directly contribute collagen to scar formation during zebrafish heart regeneration and mouse heart repair. Nat. Commun. 11, (2020).

5. Chang, J. et al. Circadian control of the secretory pathway maintains collagen homeostasis. Nat. Cell Biol. 22, 74–86 (2020).

6. Zhang, R. et al. A circadian gene expression atlas in mammals: Implications for biology and medicine. PNAS 45, 16219–16224 (2014).

7. Mure, L.S. et al. Diurnal transcriptome atlas of a primate across major neural and peripheral tissues. Science 359, (2018).

8. Hoyle, N.P. et al. Circadian actin dynamics drive rhythmic fibroblast mobilization during wound healing. Sci. Transl. Med. 9, (2017).

9. Cunningham, P. S. The circadian clock protein REVERBα inhibits pulmonary fibrosis development. PNAS 117, 1139–1147 (2020).

10. Wynn, T. A. & Vannella, K. M. Macrophages in Tissue Repair, Regeneration, and Fibrosis. Immunity 44, 450–462 (2016).

11. Witherel, C. E., Abebayehu, D., Barker, T. H. & Spiller, K. L. Macrophage and Fibroblast Interactions in Biomaterial-Mediated Fibrosis. Adv. Healthc. Mater. 8, (2019).

12. Viola, A., Munari, F., Sánchez-Rodríguez, R., Scolaro, T. & Castegna, A. The metabolic signature of macrophage responses. Front. Immunol. 10, 1–16 (2019).

13. Mantovani, A., Biswas, S. K., Galdiero, M. R., Sica, A. & Locati, M. Macrophage plasticity and polarization in tissue repair and remodelling. J. Pathol. 229, 176–185 (2013).

14. Yoo, S.-H. et al. PERIOD2::LUCIFERASE Real-Time Reporting of Circadian Dynamics Reveals Persistent *Circadian* Oscillations in Mouse Peripheral Tissues. Proc. Natl. Acad. Sci. USA 101, 5339–5346 (2004).

15. Vitaterna, M. H. et al. Mutagenesis and Mapping of a Mouse Gene, Clock, Essential for Circadian Behavior. Science 264, 719–725 (1994).

16. Yeung, C. Y. C. et al. Chick tendon fibroblast transcriptome and shape depend on whether the cell has made its own collagen matrix. Sci Rep 5, (2015).

17. Van Hoecke, L., Job, E. R., Saelens, X. & Roose, K. Bronchoalveolar lavage of murine lungs to analyze inflammatory cell infiltration. JoVE 4, (2017).

18. Naven, M. A. et al. Development of human cartilage circadian rhythm in a stem cell-chondrogenesis model. Theranostics 12, 3963–3976 (2022).

19. Calverley, B. C., Kadler, K. E. & Pickard, A. Dynamic High-Sensitivity Quantitation of Procollagen-I by Endogenous CRISPR-Cas9 NanoLuciferase Tagging. Cells 9, (2020).

20. Schwinn M.K. et al. CRISPR-Mediated Tagging of Endogenous Proteins with a Luminescent Peptide. ACS Chem. Biol. 13, 467–74 (2018).

21. Revell, C. K. et al. Modeling collagen fibril self-assembly from extracellular medium in embryonic tendon. Biophys. J. 122, 3219–3237 (2023).

22. Chen, Y., Yu, Q. & Xu, C. A convenient method for quantifying collagen fibers in atherosclerotic lesions by ImageJ software. Int. J. Clin. Exp. Med. 10, 14904–14910 (2017).

23. Martin-Burgos, B. et al. Methods for Detecting PER2:LUCIFERASE Bioluminescence Rhythms in Freely Moving Mice. J. Biol. Rhythms. 37, 78–93 (2022).

24. Yamazaki, S. & Takahashi, J. S. Real-Time Luminescence Reporting of Circadian Gene Expression in Mammals. Cell 105, 288–301 (2001).

25. Beta, R.A.A. et al. Core clock regulators in dexamethasone-treated HEK 293T cells at 4 h intervals, BMC Res. Notes 15, (2022).

26. Bhattacharya, J. & Westphalen, K. Macrophage-epithelial interactions in pulmonary alveoli. Semin. Immunopathol. 38, 461–469 (2016).

27. Chua, F. & Laurent, G. J. Fibroblasts. Encyclopedia of Respiratory Medicine (2006).

28. Sinha, M. & Lowell, C. Isolation of Highly Pure Primary Mouse Alveolar Epithelial Type II Cells by Flow Cytometric Cell Sorting. Bio. Protoc. 6, (2016).

29. Wang, N. Liang, H. & Zen, K. Molecular mechanisms that influence the macrophage m1-m2 polarization balance. Front Immunol. 28, (2014).

30. Kazlauskas A. PDGFs and their receptors. Gene 30, 1–7 (2017).

31. Bowen-Pope, D. F., Malpass, T. W., Foster, D. M. & Ross, R. Platelet-Derived Growth Factor In Vivo: Levels, Activity, and Rate of Clearance. Blood 64, 458–469 (1984).

32. Burgoyne, R. D. & Morgan, A. Secretory granule exocytosis. Physiol. Rev. 83, 581–632 (2003).

33. Gope, M. L. & Gope, R. Tyrosine phosphorylation of EGF-R and PDGF-R proteins during acute cutaneous wound healing process in mice. Wound Repair Regen. 17, 71–79 (2009).

34. Chang, J. et al. ‘Endocytic recycling is central to circadian collagen fibrillogenesis and disrupted in fibrosis’, eLife 13, (2025).

35. Shure, D., Senior, R. M., Griffin, G. L. & Deuel, T. F. PDGF AA homodimers are potent chemoattractants for fibroblasts and neutrophils, and for monocytes activated by lymphocytes or cytokines. Biochem. Biophys. Res. Commun. 186, 1510–1514 (1992).

36. Horikawa, S. et al. PDGFRα plays a crucial role in connective tissue remodeling. Scientific Reports, 5, (2015).

