## Supplementary figures for "Macrophages promote collagen deposition through circadian regulation of fibroblasts"

### Supplementary Figure 1

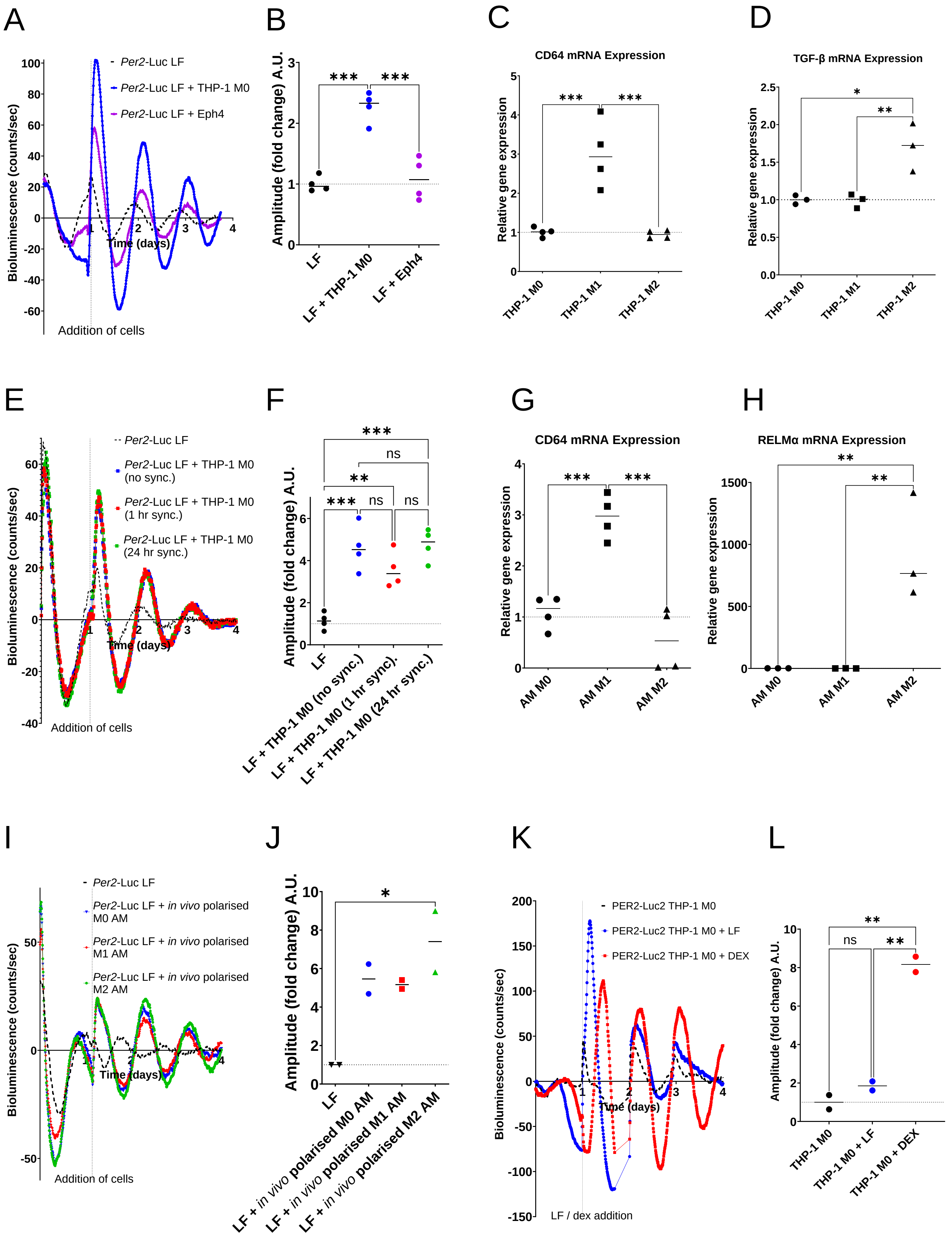

### Supplementary Figure 2

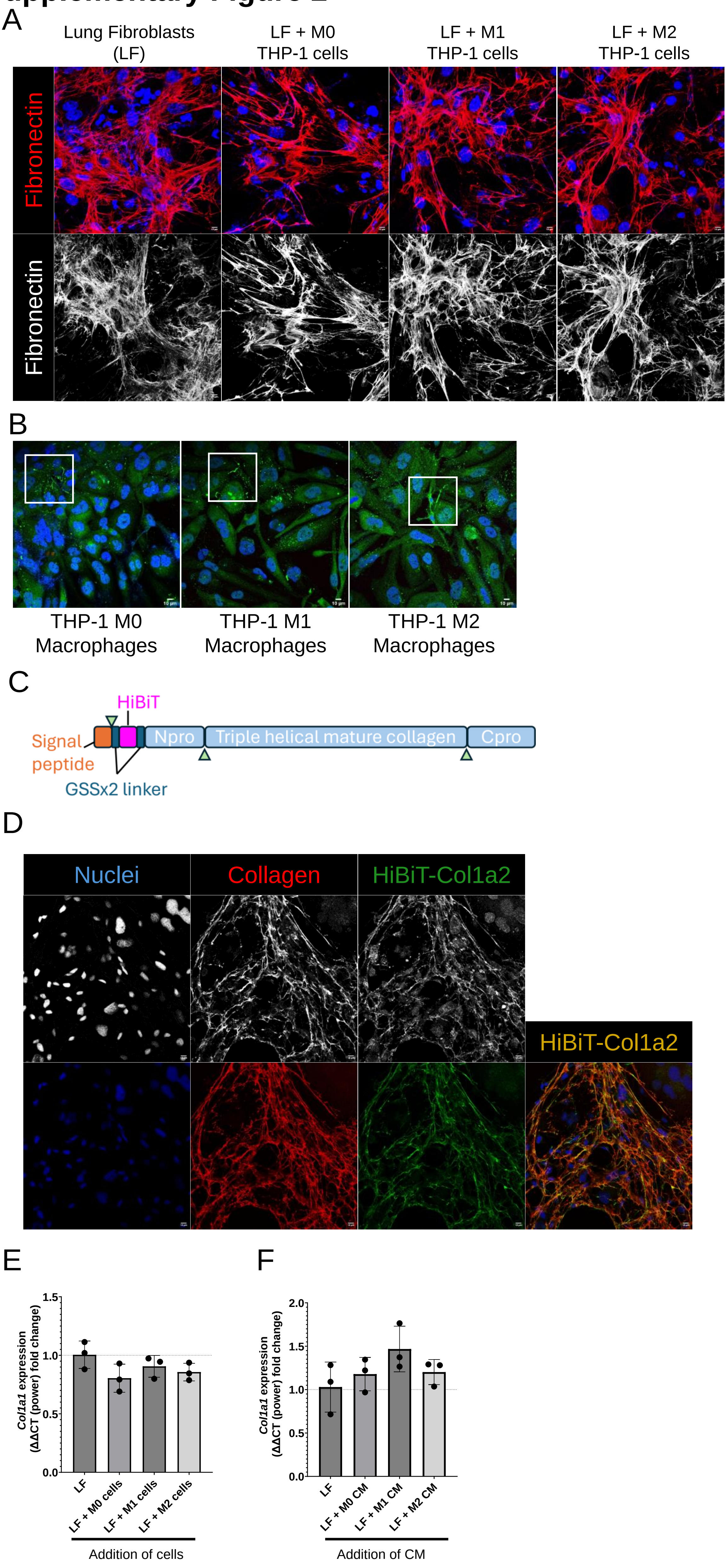

### Supplementary Figure 3

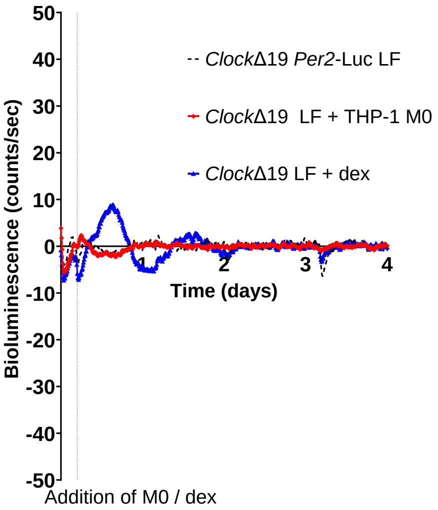

### Supplementary Figure 4

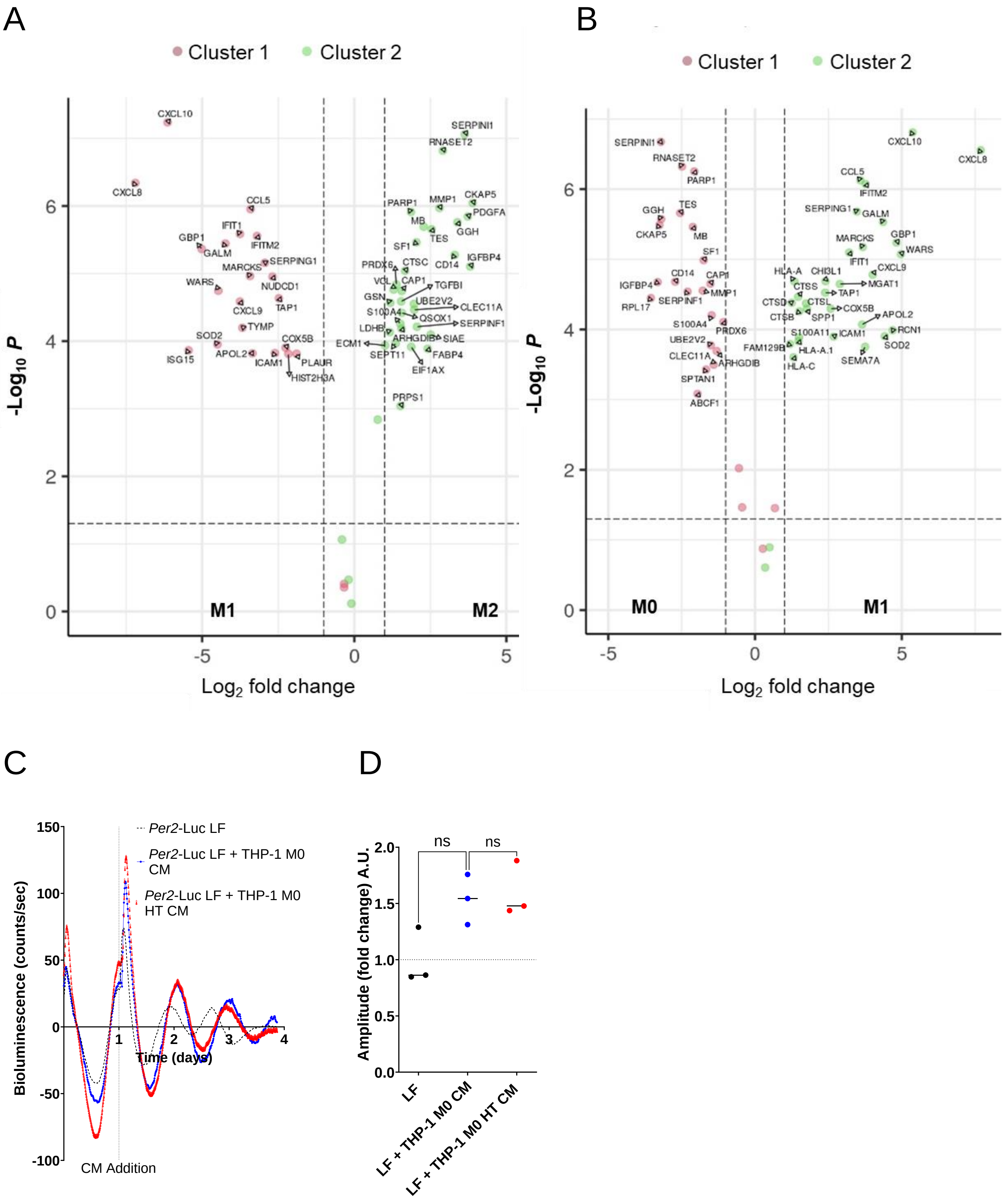

### Supplementary Figure 5

A

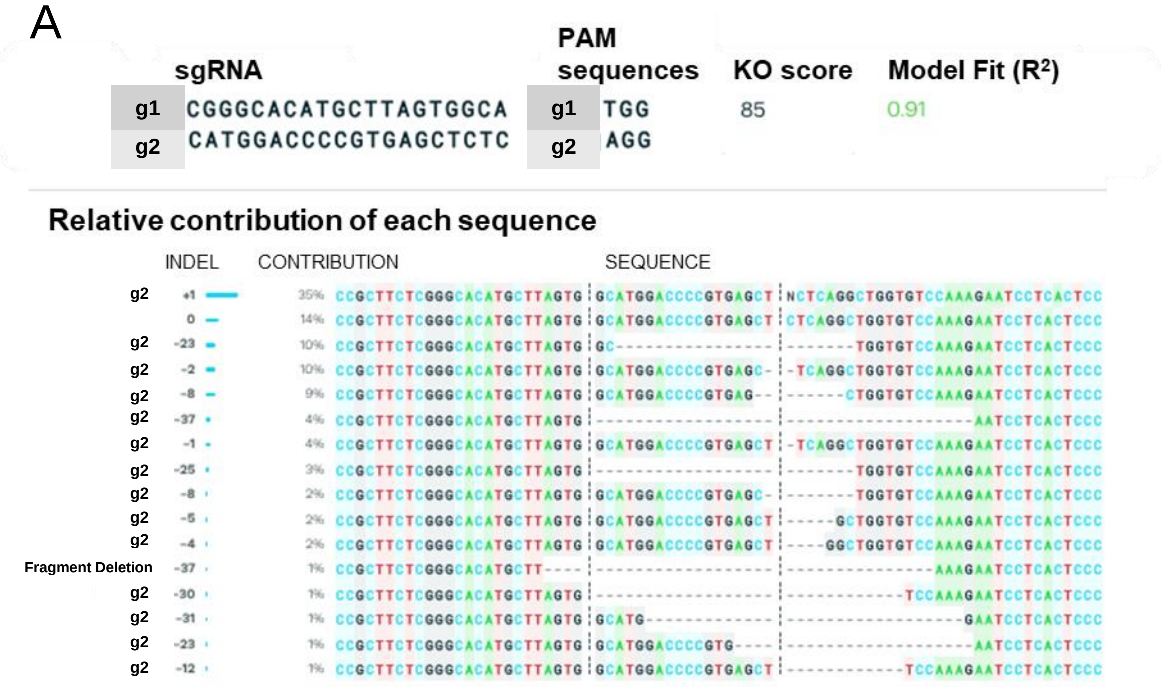


B

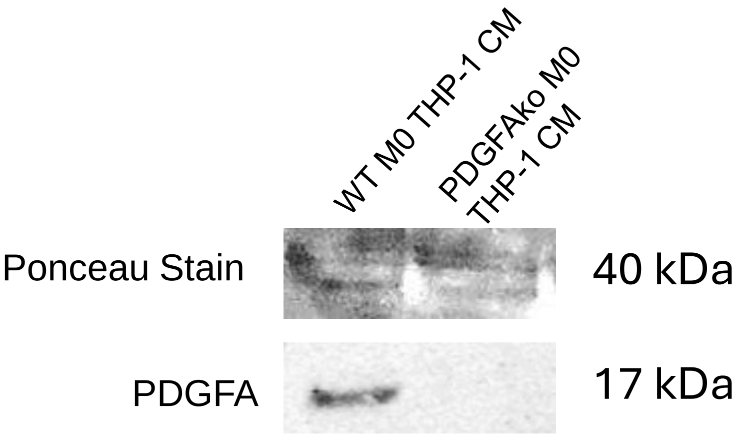
